# Modulation of cofilin 1 phosphorylation induces juvenile-like plasticity in the adult mouse visual cortex

**DOI:** 10.1101/2025.05.05.652250

**Authors:** Agustina Dapueto, Emilia Hayek, Alejo Acuña, Bruno Pannunzio, Leonel Gomez, Francesco M. Rossi

## Abstract

Cofilin 1 is an actin-depolymerizing protein that plays a fundamental role in actin dynamics, particularly within dendritic spines, where it has been implicated in both structural and functional plasticity. We recently demonstrated (by differential proteomics, western blot and immunohistochemistry) that the expression of cofilin 1 and its inactive phosphorylated form are dynamically regulated in the mouse visual cortex during postnatal development and by visual experience. Moreover, we found that cofilin 1 expression levels correlates with periods of heightened plasticity in the mouse visual cortex.

In this study, we analyzed whether the pharmacological modulation of cofilin 1 activity affects plasticity processes in the visual cortex. Adult mice were treated with a synthetic peptide inhibitor of cofilin 1 activity (PCOF) or a control peptide (TAT) and then monocularly deprived of vision during 3 days. Following reopening of the deprived eye, structural plasticity was assessed by quantifying dendritic spine density using Golgi-like staining, and visual plasticity was evaluated by measuring visual acuity through the optomotor response test.

Our results show that, in adult mice treated with the PCOF peptide - but not in controls - monocular deprivation led to a significant reduction in dendritic spine density in the contralateral visual cortex, as well as a decrease in visual acuity of the previously deprived eye. These findings indicate that cofilin 1 activity is crucial for the regulation of experience-dependent plasticity in the adult mouse visual cortex.

## Introduction

The visual cortex is a paradigmatic model for studying experience-dependent plasticity. Several studies have shown that manipulating the visual input by a simple occlusion of vision in one eye for a few days (monocular deprivation, MD) induces dramatic and permanent changes in the visual cortex at both structural and functional level, as well as loss of vision. These effects are quite marked when MD is done during a particularly sensitive or critical period of postnatal life (CP), while they are reduced or absent when MD is done during adulthood (Hensch, 2004; Levelt & Hubener, 2012). Over the past two decades, several studies have demonstrated that juvenile-like plasticity can be reinstated in adults through various treatments. Prominent examples of this phenomenon are, for instance, chronic fluoxetine administration, rearing in enriched environment, or feeding animals with caloric restriction (Hensch et al., 1998; Fagiolini & Hensch, 2000; Pizzorusso et al., 2002). This discovery led to the successful employment of such treatments to facilitate recovery from visual pathology (e.g.: amblyopia) (Sale et al., 2007; Maya-Vetencourt et al., 2008; Spolidoro et al., 2011; Baroncelli et al., 2012).

The cellular and molecular mechanisms underlying the opening, duration and closure of the critical period and the processes controlling plasticity levels are still largely unknown. However, various studies pointed out that the development of the excitatory/inhibitory tone and the maturation of structural brakes in the visual cortex are fundamental players (Hensch, 2005; Tropea et al., 2009; Fawcett et al., 2019). Large-scale studies (at both transcriptomic and proteomic levels) have suggested that a substantial number of genes regulate plasticity and that experience leaves distinct “molecular fingerprints” that vary according to the degree of cortical plasticity (see Rossi & Pizzorusso, 2025 for a review). Our group recently used a proteomic approach to identify protein profiles differentiating the visual cortex during the critical period (characterized by high plasticity) from adulthood (characterized by low plasticity), and following fluoxetine treatment in adults (characterized by pharmacologically-induced high plasticity) (Ruiz-Perera, et al., 2015; Bornia et al., 2020). Among various differentially expressed proteins, cofilin 1 emerged as a relevant one as its levels were high, in correlation with high plasticity (critical period and fluoxetine treatment), and low, in correlation with low plasticity (adults). We further demonstrated by western blot and inmunohistochemistry that cofilin 1 and its inactivated form (phosphorylated in Ser3), are dynamically modulated during postnatal development and that MD affects their expression during the CP but not in adults (Laguardia et al., 2023). These results suggested for the first time cofilin 1 as a potential player in the control of visual cortex plasticity.

In the present work, we administered an inhibitory peptide targeting cofilin 1 activity to adult mice in vivo, and analyzed MD-induced effects in the visual cortex as marker of experience-dependent plasticity. Specifically, we investigated MD effects on structural parameters (dendritic spine density, using the Golgi-like staining), and on visual parameters (spatial resolution threshold, using the optomotor test). Our data indicate that cofilin 1 activity modulation reinstates high plasticity levels in the adult mouse visual cortex.

## Results

### Analysis of dendritic spine density in the mouse visual cortex with the Golgi-like staining

We used the commercial *elite*Golgi kit (Bioenno) to stain and analyze spine density on apical dendrites of LII-III and LV pyramidal neurons of the binocular visual cortex. Following the fine-tuning of the methodology, we analyzed spine density in the visual cortex of untreated CP and adult mice (Supplementary Figure 1). We observed that spine density (n protrusions *100μm) in the visual cortex is higher in young mice than in adults, decreasing approx. 55% from CP to adulthood (CP, P28: 176.2±3.4, n=24 dendrites, n=5 mice, vs. AD, P58: 79.4±1.6, n=32 dendrites, n=5 mice, Student’s t-test, p<0.05).

In order to confirm that the methodology in use allows the detection of plasticity at structural level, we compared the effects of MD on dendritic spine density between critical period and adult mice. CP and adult mice were deprived of vision in one eye (right eye) by lid suturing during 3 days from P28 to P31 and from P59 to P62, respectively. Then, mice were sacrificed and brains processed for Golgi-like staining (Supplementary Figure 2). We observed that during the critical period, MD reduces spine density in the contralateral vs. ipsilateral visual cortex of approximately 36% (contral.: 113.9±1.5 n=28 dendrites, vs. ipsil.: 178.0±3.7, n=24 dendrites, n=6 mice, Student’s t-test, p<0.05). On the contrary, no effects of MD were observed in adult mice (contral.: 86.4±3.1 n=24 dendrites, vs. ipsil.: 87.2±3.6, n=21 dendrites, n=6 mice, Student’s t-test, p=0,88), confirming that adult mice are less plastic than young ones.

### Modulation of cofilin activity reinstates experience-dependent structural plasticity in the adult mouse visual cortex

Finally, in order to analyze if cofilin 1 plays a role in structural plasticity of the mouse visual cortex we made use of the pharmacological modulation of its activity by intravenous administration of a single dose of the PCOF inhibitory peptide or the TAT control peptide at a concentration of 15 pmol/g. First we checked if peptides treatment *per se* affected spine density in normal adult mice treated at P58 and analyzed 4 days later at P62 (Figure 1). We observed no differences in spine density between the ispilateral (right) and the contralateral (left) visual cortex of adult mice treated with the TAT peptide (contral.: 82.2±1.9 n=19 dendrites vs. ipsil.: 81.0±1.4 n=19 dendrites, n=8 mice, Student’s t-test, p=0.60) or with the PCOF peptide (contral.: 84.2±1.7 n=19 dendrites vs. ipsil.: 83.0±1.5 n=19 dendrites, n=8 mice, Student’s t-test, p=0.63).

**Figure 1.**
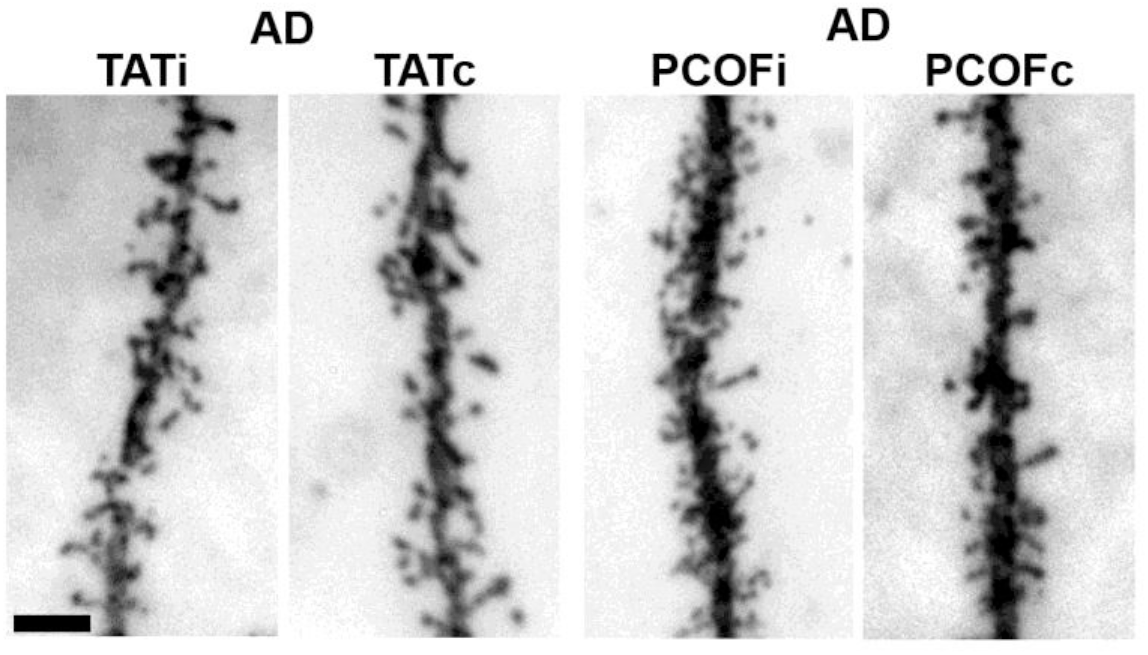
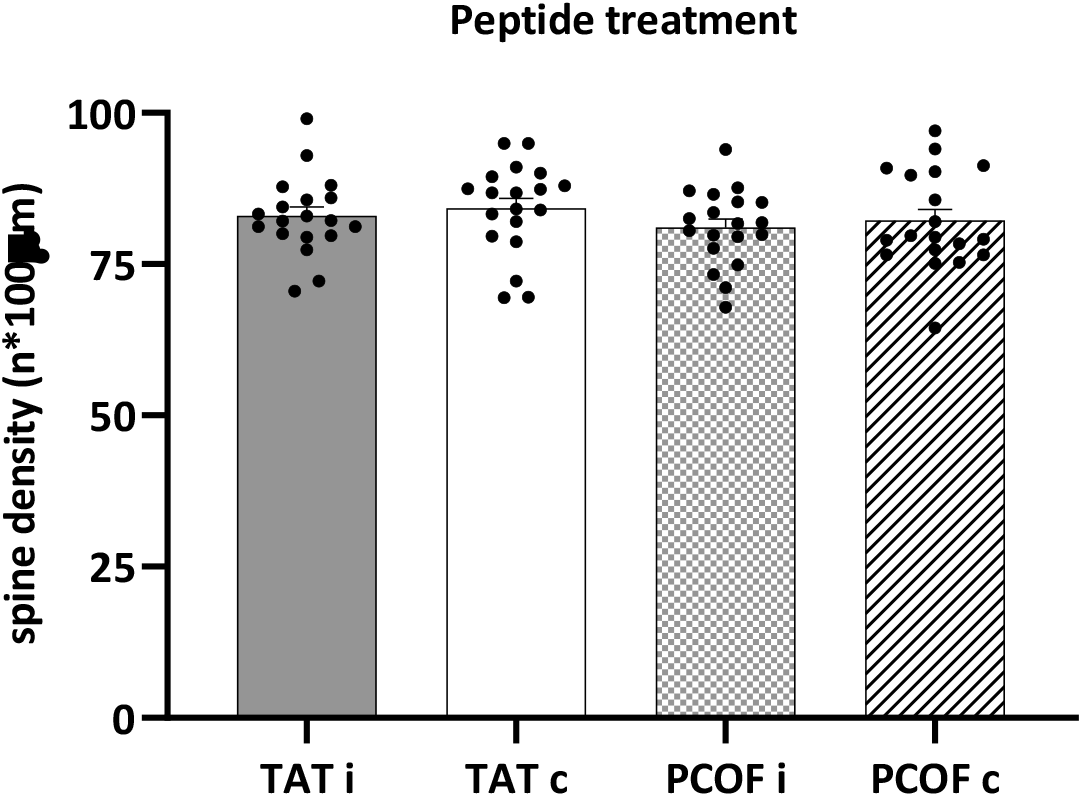
Spine density is not affected by peptides administration in normal adult mice. Representative images of apical dendrites from the ispilateral and contralateral visual cortex of TAT treated (TATi and TATc) and PCOF treated (PCOFi and PCOFc) adult (AD) mice. Bar: 5 μm. The scatter plot represents the spine density (n protrusions*100 μm) in the ispilateral (TATi, right) and contralateral (TATc, left) visual cortex of n=8 adult mice treated with the TAT peptide, and in the ispilateral (PCOFi, right VC) and contralateral (PCOFc, left VC) visual cortex of n=8 adult mice treated with the PCOF peptide. Dots represent spine density values of individual dendrites; mean is marked as a bar±SEM. Mean spine density: TATi: 81.0±1.4 n=19 dendrites vs. TATc: 82.2±1.9 n=19 dendrites, n=8 mice, Student’s t-test, p=0.60; PCOFi: 83.0±1.5 n=19 dendrites vs. PCOFc: 84.2±1.7 n=19 dendrites, n=8 mice, Student’s t-test, p=0.63.

Interestingly, we observed that treating adult mice with the PCOF peptide reinstates the juvenile-like MD induced effects at structural level. One day following PCOF treatment (P58), 3 days monocular deprivation (P59-62) reduces spine density in the contralateral vs. ipsilateral visual cortex of adult mice of approximately 42% (contral.: 49.4±1.6 n=37 dendrites vs. ipsil.: 84.9±2.5 n=37 dendrites, n=8 mice, Student’s t-test, p<0.05). On the contrary, no effects were observed in deprived adult mice treated with the control TAT peptide (contral.: 84.1±2.1 n=33 dendrites vs. ipsil.: 81.6±2.3 n=33 dendrites, n=8 mice, Student’s t-test, p=0,69) (Figure 2).

**Figure 2.**
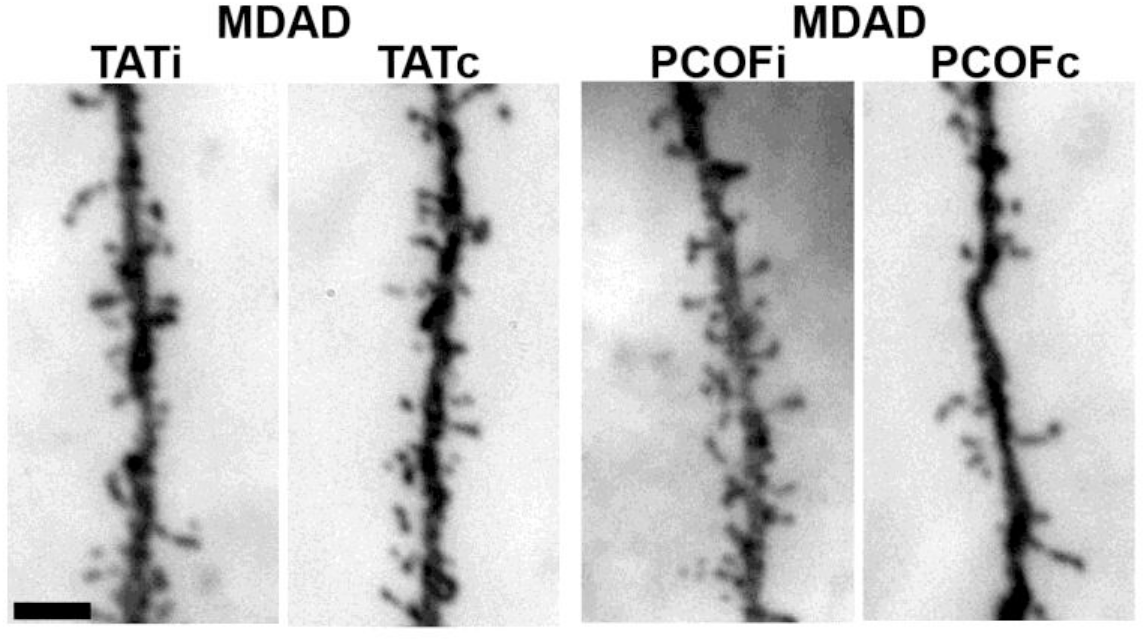
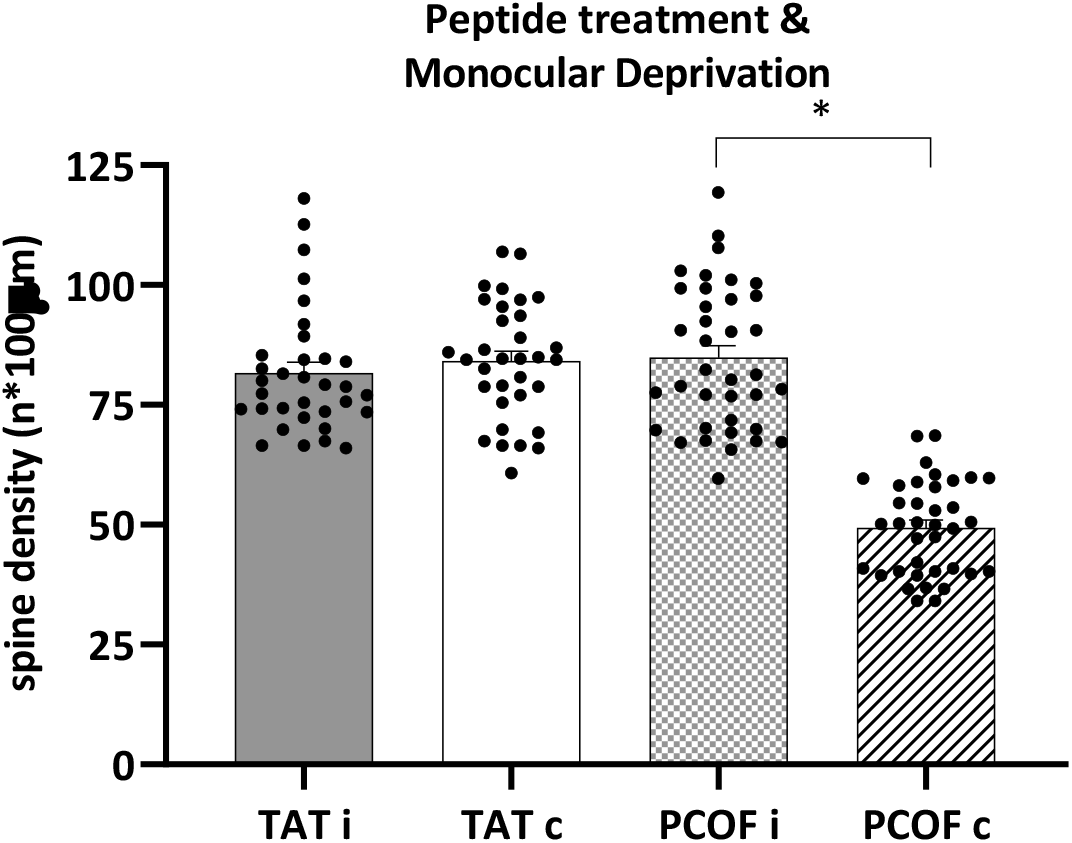
Monocular deprivation reduces spine density in the contralateral visual cortex in PCOF treated mice, but not in TAT treated ones. Representative images of apical dendrites from the ispilateral and contralateral visual cortex of TAT treated (TATi and TATc) and PCOF treated (PCOFi and PCOFc) deprived (MD, right eye) adult (AD) mice. Bar: 5 μm. The scatter plot represents the spine density (n protrusions*100 μm) in the ispilateral (TATi) and contralateral (TATc) visual cortex of n=8 deprived adult mice treated with the TAT peptide, and in the ispilateral (PCOFi) and contralateral (PCOFc) visual cortex of n=8 deprived adult mice treated with the PCOF peptide. Dots represent spine density values of individual dendrites; mean is marked as a bar±SEM. Mean spine density: TATi: 81.6±2.3 n=33 dendrites vs. TATc: 84.1±2.1 n=33 dendrites, n=8 mice, Student’s t-test, p=0.69; PCOFi: 84.9±2.5 n=37 dendrites vs. PCOFc: 49.4±1.6 n=37 dendrites, n=8 mice, Student’s t-test, *p<0.05.

### Analysis of spatial resolution threshold with the optomotor test

We used the optomotor test to measure spatial resolution threshold of the left and right eye in unrestrained mice as an index of visual acuity (Douglas et al., 2005). Following the fine-tuning of the methodology, we analyzed spatial resolution threshold (c/d) in CP and adult mice (Supplementary Figure 3). In agreement with previous studies (Prusky et al., 2004), we observed that spatial resolution threshold is similar in critical period and adult mice (CP, P28: 0.37±0.01, n=6 mice, vs. AD, P60: 0.38±0.01, n=10 mice; Student’s t-test, p=0.85).

In order to confirm that the optomotor test allows detecting experience-dependent plasticity changes of visual parameters, we verified if MD induces a modification of spatial resolution threshold comparing critical period and adult mice (Supplementary Figure 4). CP and adult mice were deprived of vision by lid suturing during 3 days from P28 to P31 and from P59 to P62, respectively, and then analyzed in the optomotor test. We found that during the critical period, MD reduces spatial resolution threshold in the deprived eye vs. the fellow eye of approximately 43% (fellow eye: 0.38±0.02 vs. MD eye: 0.20±0.01, n=6 mice, Student’s t-test, p<0.05). On the contrary, no effects of MD were observed in adult mice (fellow eye 0.39±0.01 vs. MD eye: 0.38±0.01, n=6 mice, Student’s t-test, p=0.81).

### Modulation of cofilin activity reinstates experience-dependent visual plasticity in the adult mouse visual cortex

Finally, to analyze if cofilin 1 plays a role on the MD-induced effects on visual acuity of the mouse visual cortex we used the administration of the PCOF or the TAT peptide and analyzed spatial resolution threshold. We first checked the effects of peptides treatment in normal adult mice (treated at P58, analyzed at P62) (Figure 3). We observed no differences on spatial resolution threshold between the two eyes in adult mice treated with the PCOF peptide (left eye: 0.39±0.01 vs. right eye: 0.39±0.01, n=6 mice, Student’s t-test, p=0.91) nor in TAT control-treated mice (left eye: 0.38±0.01 vs. right eye: 0.41±0.01, n=6 mice, Student’s t-test, p=0.89).

**Figure 3.**
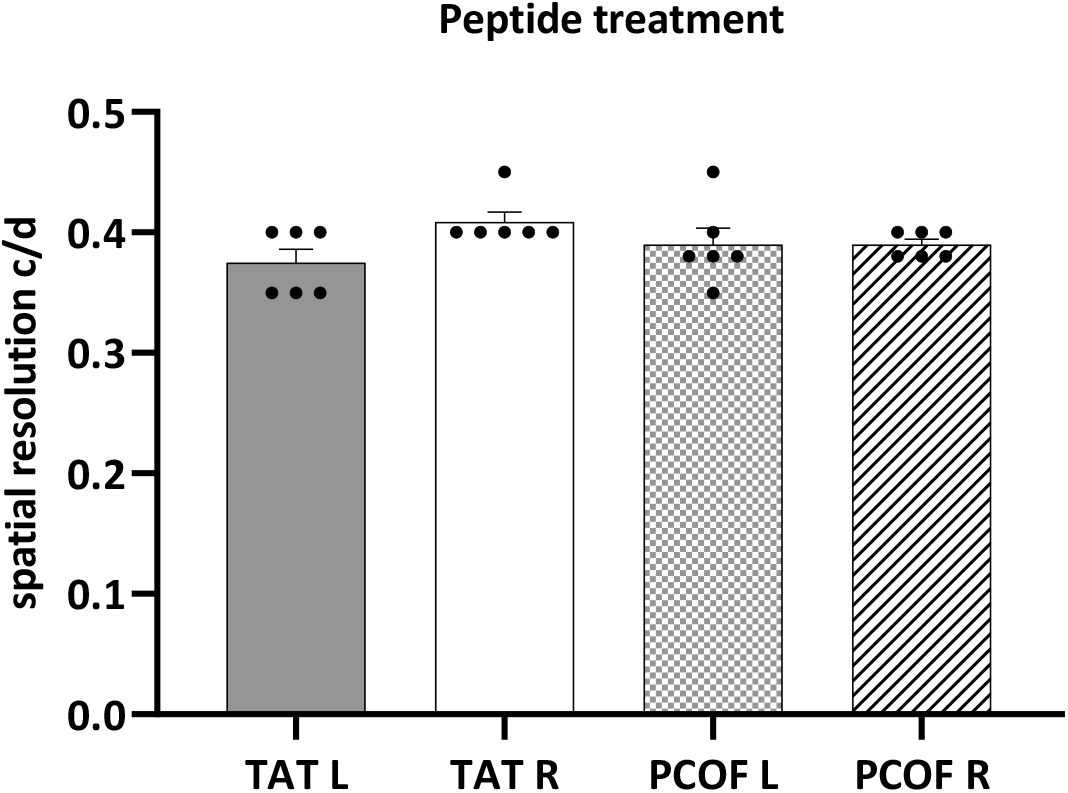
Visual acuity is not affected by peptides administration in normal adult mice. The scatter plot represents the spatial resolution (c/d) through the left eye (TAT L) and the right eye (TAT R) of n=6 adult mice treated with the TAT peptide, and through the left eye (PCOF L) and the right eye (PCOF R) of n=6 adult mice treated with the PCOF peptide. Dots represent VA values of individual mice; mean is marked as a bar±SEM. Mean VA: TAT L: 0.38±0.01 vs. TAT R: 0.41±0.01, n=6, Student’s t-test, p=0.89; PCOF L: 0.39±0.01 vs. PCOF R: 0.39±0.01, n=6, Student’s t-test, p=0.91.

On the contrary, we found that one day following PCOF peptide treatment (P58), 3 days monocular deprivation (P59-62) reduces spatial resolution threshold in the deprived eye vs. the fellow eye of approximately 31% (fellow eye: 0.38±0.01 vs. MD eye: 0.27±0.01, n=16 mice, Student’s t-test, p<0.05), while no effects were observed in deprived adult mice treated with the TAT control peptide (fellow eye: 0.39±0.01 vs. MD eye: 0.39±0.01, n=12 mice, Student’s t-test, p=0.81) (Figure 4).

**Figure 4.**
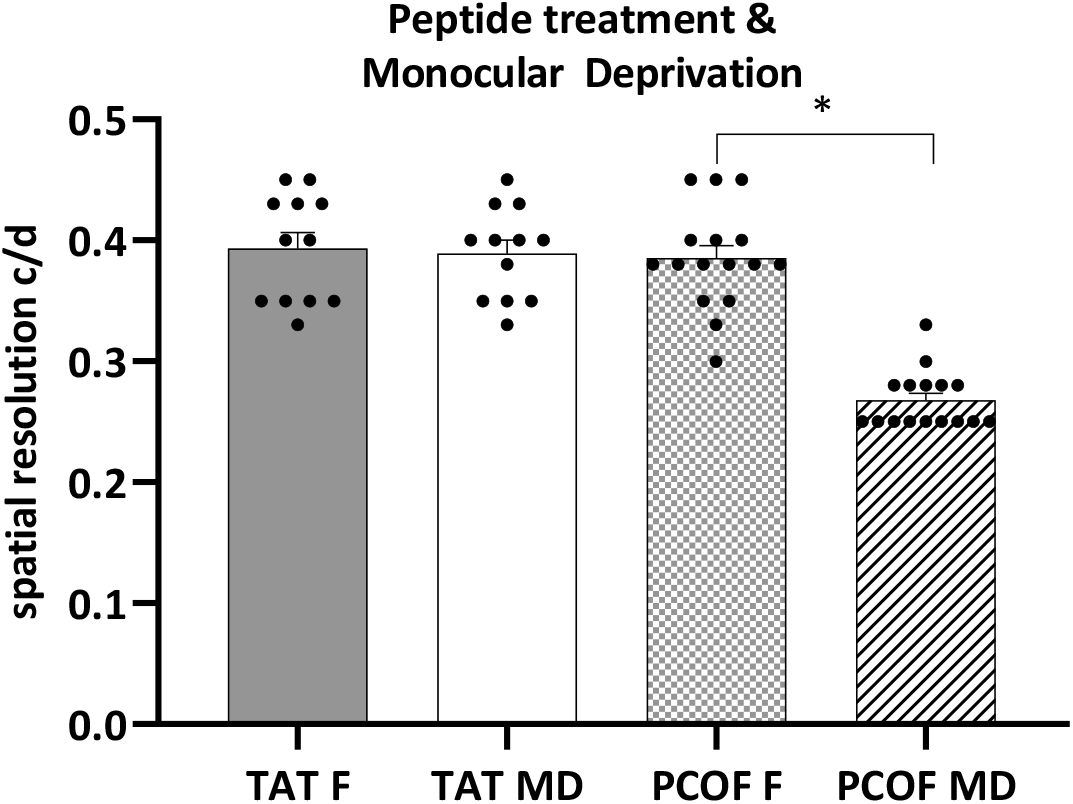
Monocular deprivation reduces visual acuity through the deprived eye in PCOF treated mice, but not in TAT treated ones. The scatter plot represents the spatial resolution (c/d) through the fellow eye (TAT F) and the deprived eye (TAT MD) of n=12 adult (AD) mice treated with the TAT peptide, and through the fellow eye (PCOF F) and the deprived eye (PCOF MD) of n=16 adult mice treated with the PCOF peptide. Dots represent VA values of individual mice; mean is marked as a bar±SEM. Mean VA: TAT F: 0.39±0.01 vs. TAT MD: 0.39±0.01, n=12, Student’s t-test, p=0.81; PCOF F: 0.38±0.01 vs. PCOF MD: 0.27±0.01, n=16, Student’s t-test, *p<0.05.

## Materials and Methods

### Animals and treatment

All experiments were performed in accordance with the EC Directive 86/609/EEC for animal experiments (directive from the European Commission of the European Economic Community) and with the Uruguayan Research Ethic Committees (protocol approval #1131, ExpFCien 2400 11-000556-20). A total of 74 C57BL6/J mice (both sexes) at different postnatal (P) ages (P28, n=16, P58: n=58) provided by the Unidad de Reactivos y Biomodelos de Experimentación, Facultad de Medicina, Universidad de la República, Montevideo, Uruguay, were used. Mice were bred under specific pathogen-free conditions, housed (6/cage) in ventilated cages with positive pressure (20±1°C, relative humidity 40–60 %, 14/10 h. light-dark cycle), fed with standard mouse diet *ad libitum*, and had free access to water.

### Monocular deprivation

Monocular deprivation was performed during 3 days in critical period (from P28 to P31) or in adult (from P58/59 to P61/62) mice anesthetized with ketamine / xylazine / acedan 100 / 10 / 3 mg/Kg, by suturing the eyelids of the right eye as previously described (Laguardia et al., 2023). The integrity of the suture was checked daily and mice were used only if the eyelids remained closed throughout the duration of the deprivation period. The eyelids were reopened immediately before performing the optomotor test and the pupil was checked for clarity. Then mice were sacrificed (by cervical dislocation) and brain tissues collected. A total of 40 mice (P28: n=6, P58/59: n=34) was monocularly deprived of vision.

### Peptide treatment

In order to modulate cofilin 1 phosphorylation *in vivo* we used a peptide (GenScript) consisting of the first 16 amino acids of cofilin 1 with the residue Ser3 phosphorylated (Duffney et al., 2015). The phospho-cofilin peptide was coupled to the protein transduction domain of the HIV TAT protein to make it cell permeable (PCOF peptide: RKKRRQRRRMA{pSER}GVAVSDGVIKVFN). As a control, the peptide containing only the TAT-HIV domain was used (TAT peptide: RKKRRQRRR). Peptides were re-suspended in sterile water at a stock concentration of 2.5 mM, diluted in physiological solution at 5 microM (5 pmol/microL) and delivered by an intravenous injection (i.v.) at 15 pmol/g. A total of 44 mice (TAT: n=20, PCOF: n=24) was injected with peptides.

### Golgi staining

Mice were sacrificed by cervical dislocation, brains were quickly removed, rinsed in physiological solution and were Golgi-stained using the *élite* Golgi Kit (#006690, Bioenno, Tech, LLC). Briefly, brains were immersed in the impregnation solution and stored at RT in darkness for 6 days. Brains were cut with a vibratome (LeicaGeoSystems VT1000S) in 150 *μ*m thick coronal sections that were collected in PB 0.1M (left and right visual cortex were separately collected). Sections were incubated free-floating in the staining and clarity solutions, rinsed in PBST 0.01M and directly mounted on gelatin-coated slides. Following dehydration (ethanol 95% and 100%) and cleaning (xylene), slides were covered with coverslips in mounting media (Canadian balsam, Biopack, or Entellan, Sigma-Aldrich). A total of 54 mice (CP: n=11, AD: n=43) was processed for Golgi staining.

### Spine density analysis

Photographs of the images were obtained using a D70S Digital Camera (Nikon) attached to a Microphot-FXA microscope (Research Microscope, Nikon) with a 40x objective. Spine density analysis was carried out on apical dendrites of pyramidal neurons in layers II-III and V of the binocular visual cortex (bVC). bVC position was identified comparing sections with images from the mouse brain atlas (Paxinos & Franklin, 2007; AP: −4.36 and −3.08; ML: 3.25 and 2.5). Nissl-stained alternating sections were used to identify cortical layers. Only dendrites of neurons showing typical pyramidal neuron morphology were analyzed (Braak & Braak, 1985; Spruston, 2008). Furthermore, in order to be analyzed dendrites must meet the following criteria: (i) to be viewed clearly, without significant breaks or interrupted staining; (ii) not to be interrupted by others stained neurons; (iii) the soma must be identifiable with clarity and be completely stained. Once the dendrites selected, the number of protrusions per 100 μm segment, along the entire dendrite was counted. Image analysis was carried out with the free software Fiji (Fiji Is Just ImageJ, NIH).

### Optomotor test

Behavioral threshold acuity was evaluated using the optomotor task in unrestrained mice (Prusky et al., 2004; Douglas et al., 2005). A virtual-reality chamber was created with four monitors facing into a square, a mirror in the floor and another in the roof of the chamber (with a small hole in its center were the camera was positioned). A platform was secured on the bottom of the floor (height 13 cm, diameter 5 cm). The stimulus (vertical sine-wave gratings) was projected on the monitors using the PsychoPy® software (PsychoPy v3.0, www.psychopy.org). The stimulus changed in spatial frequency (0.1, 0.2., 0.3, 0.35, 0.375, 0.4, 0.425, 0.45 c/d; duration: 60 sec. each) and direction (alternating clockwise and counterclockwise, each 30 sec.) and was projected at 12 deg/sec. velocity and 100% contrast. The experiments were performed in scarce luminance room conditions (approx. 12 cd/m^2^). All animals were habituated before the onset of testing by gentle handling and by placing them on the arena platform for a few minutes at a time. Experimenters were blind to treatments conditions of the mice and the animal’s previously recorded thresholds. Videos were analyzed counting number of head movements and comparing with the direction and frequency of the stimulus. Maximal spatial resolution visual threshold was taken as the last frequency at which mice responded in agreement with the direction of the stimulus. A total of 70 mice (CP: n=16, AD: n=54) was processed for the optomotor test.

## Discussion

The present study aimed at investigating whether cofilin 1 plays a role in experience-dependent plasticity.

Cofilin 1 is an actin-depolymerizing protein belonging to the ADF/cofilin family, which in mammals comprises three members: cofilin 1 (non-muscle, or n-cofilin), cofilin 2 (muscle-specific, or m-cofilin), and ADF (actin depolymerizing factor, also known as destrin). Among these, cofilin 1 and ADF are highly expressed in the brain, particularly at excitatory synapses (Racz and Weinberg, 2006; Bellenchi et al., 2007; Rust et al., 2010; Görlich et al., 2011). The activity of ADF/cofilin is tightly regulated by phosphorylation at serine residue 3 (Ser3), which inhibits their binding to actin filaments - an important post-translational modification controlling their function (Yang et al., 1998; Meng et al., 2002). Cofilin 1 plays a crucial role in actin filament turnover, especially within dendritic spines, where it has been extensively linked-primarily through studies in the hippocampus - to both structural and functional synaptic plasticity (Rust, 2015; Bamburg et al., 2021; Namme et al., 2021). Different experimental strategies have been used to modulate cofilin 1 expression (knock out, conditional transgenic mutants, or interference RNA) or its activity (phospho-mimetic, or peptides administration) (Shaw & Bamburg, 2017). Here we used the administration of a synthetic phosphorylated peptide that competes with the endogenous phosphorylated cofilin as a substrate for the slingshot phosphatase (SSH1) responsible for cofilin 1 dephosphorylation, thus determining a reduction in its activity (Shaw & Bamburg, 2017). Previous works have shown that *in vivo* treatment with these peptides effectively induce a modulation of the phosphorylation state of cofilin 1 (Aizawa et al., 2001; Toda et al., 2006; Wang et al., 2013; Zhou et al., 2014; Duffney et al., 2015; Rust, 2015; Tsubota et al., 2015; Deng et al., 2016).

In order to study brain plasticity processes, we exploited the solid model of monocular deprivation to study structural and functional effects in the adult mouse visual cortex. We studied MD-effects on dendritic spine density as structural parameter and on visual acuity as physiological parameter.

Initially, we confirmed that dendritic spine density decreases from the critical period to adulthood which is in agreement with previous studies and it is possibly related to the end of the postnatal pruning process (Runge et al., 2020). We then verified if the experimental approach was able to detect plasticity at structural level and we found that MD decreases spine density when done in critical period mice but not in adults. These results confirmed previous works (Mataga et al., 2004; Bochner et al., 2014; Bukhari et al., 2015; see also Djurisic et al., 2003; Hofer et al., 2009) and allowed the use of this experimental strategy to detect the potential effect of cofilin 1 on visual cortex plasticity at structural level. Finally we analyzed whether *in vivo* modulation of cofilin 1 activity restores sensitivity to MD in the adult visual cortex. We found that spine density decreases in the contralateral visual cortex of monocularly deprived adult mice treated with the PCOF peptide, which is the expected result for a treatment with a potential plasticity-increasing effect. A similar MD-induced decrease in cortical spine density has been observed by others for instance in adult Lynx1 (Bukari et al., 2015) and PSD95 (Yusifov et al., 2021) knockout mice, demonstrating the role of these factors in cortical plasticity. Additionally, it was shown that sleep alterations can alter the elimination/formation ratio in monocularly deprived adult mice (Zhou, et al., 2020).

As for visual function, we used the optomotor test to detect spatial resolution threshold which is considered an index of visual acuity. While the optomotor response is a behavioral test reflecting sub-cortical and cortical function (Douglas et al., 2005), the results obtained with such test have been shown to be consistent with electrophysiological cortical measures (such as visual evoked potentials) and to respond to experimental manipulations similarly (Durand et al. 2012; Kang et al., 2013). We then showed that interfering with cofilin 1 activity decreases visual acuity in the deprived eye of adult mice.

Besides electrophysiological VEP measures, other behavioral tests have been used to detect restored plasticity in adult rodents or recovery of vision in adult amblyopic animals. For example, it has been shown that fluoxetine or sub anesthetic ketamine administration, as well as enriched environment facilitate the recovery of normal visual acuity values (measured with the visual water maze task) in the deprived eye of amblyopic adult mice (Sale et al., 2007; Maya-Vetencourt et al., 2008; Grieco et al., 2020).

The present data indicate that cofilin 1 is a relevant player in the control of experience-plasticity in the mouse visual cortex as its modulation *in vivo* favors reopening of high plasticity levels at adult stages. These results are in agreement with previous works indicating a similar role of cofilin 1 in other brain areas or plasticity processes. It has been shown that cofilin 1 is important in hippocampal long-term depression/potentitation (Cao et al., 2017), in transcolumnar plasticity of the somatosensory cortex (Tsubota et al., 2015), as well as in memory processes (Medina et al., 2020) (for reviews see: Rust, 2015; Shaw & Bamburg, 2017; Namme et al., 2021). Particularly lined-up with the present results is the study by Cerri et al., 2011. In this work the authors used the bacterial protein toxin, cytotoxic necrotizing factor 1 (CNF1), to induce a persistent activation of Rho GTPases. They showed that *in vivo* administration of CNF1 reinstates ocular dominance plasticity in the adult rat visual cortex. Rho GTPases, mainly Rac1, Cdc42, and RhoA, are proteins that regulate actin activity. Interestingly, cofilin 1 activity is one of the effectors of RhoGTPases: Rac1 can activate LIMK1, while Rho and Cdc42 can activate LIMK2, the two main enzymes responsible for the phosphorylation, i.e. inactivation, of cofilin 1 (Sumi et al., 1999; Name et al., 2021). One possible interpretation of these findings is that the CNF1-induced restoration of high plasticity levels in adult mice is partially mediated by reduced cofilin 1 activity, reflected by an increase in its phosphorylated (inactive) form. This interpretation is supported by our results showing that administration of the PCOF peptide - which enhances endogenous phosphocofilin levels by inhibiting the corresponding phosphatase - promotes cortical plasticity.

In summary, our findings highlight a critical role for cofilin 1 activity in regulating experience-dependent plasticity in the mouse visual cortex. However, the precise molecular mechanisms by which cofilin 1 exerts these effects remain to be elucidated.

## Supporting information

Dapueto et al 2025 Suppl revised

## Acknowledgements

This work was supported by Programa de Desarrollo de las Ciencias Básicas (Pedeciba), Agencia Nacional de Investigación a Innovación (ANII), and Comisión Sectorial de Investigación Científica (CSIC). The authors wish to thank Dr. F. Zolessi, Dr. L. Gomez and Dr. A. Villarino (Facultad de Ciencias, Universidad de la República, Montevideo, Uruguay) for experimental support and assistance, and for providing comments on the manuscript; Dr. R. Mazziotti, L. Baroncelli and T. Pizzorusso (Scuola Normale Superiore/Institute of Neuroscience, CNR, Pisa, Italy) for assisting in the setting-up of the optomotor response test. F. M. Rossi: Conceptualization, Methodology, Software, Data curation, Writing, Original draft preparation, Supervision, Reviewing and Editing, Funding acquisition; A. Dapueto, L. Gomez, E. Hayek, A. Acuña, B. Pannunzio: Investigation, Visualization. A. Dapueto was supported by a CSIC grant (#567, Proyecto de Iniciación a la Investigación). The authors confirm that the funders had no influence over the study design, content of the article, or selection of this journal.

