## Supplementary material for "Modulation of cofilin 1 phosphorylation induces juvenile-like plasticity in the adult mouse visual cortex": Dapueto et al 2025 Suppl revised

#### Supplementary Figure 1

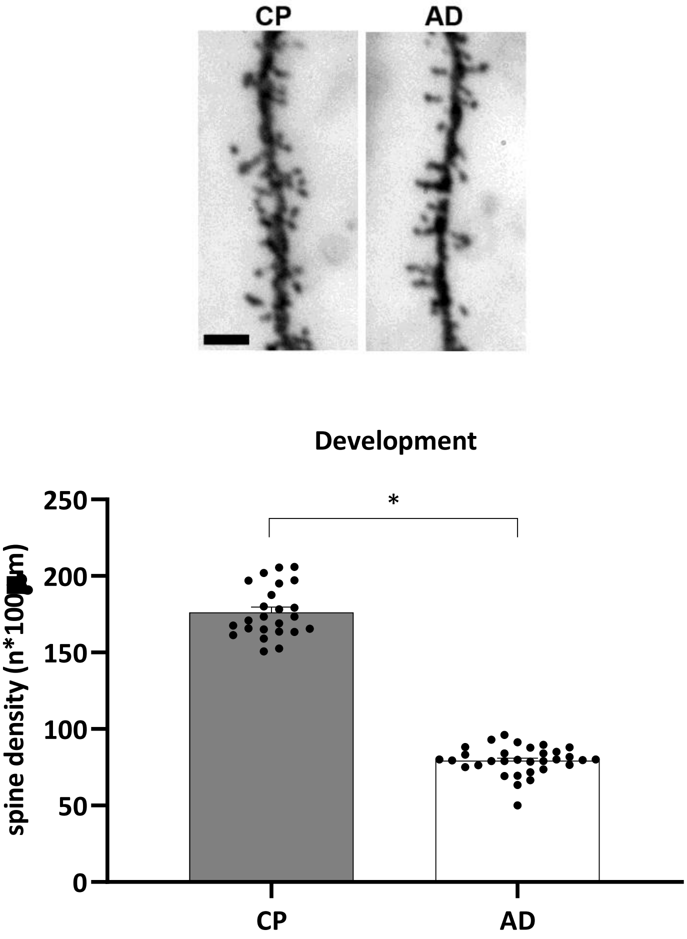


**Supplementary Figure 1:** Spine density decreases from critical period to adulthood in the visual cortex of mice. Representative images of apical dendrites from the visual cortex of critical period (CP) and adult (AD) mice. Bar: 5 µm. The scatter plot represents the spine density (n protrusions*100 µm) in the visual cortex of n=5 CP and n=5 AD mice. Spine density of the right and left VC were pooled. Black dots represent spine density values of individual dendrite; mean is marked as a bar±SEM. Mean spine density: CP: 176.2±3.4, n=24 dendrites, n=5 mice vs. AD: 79.4±1.6, n=32 dendrites, n=5 mice, Student´s t-test, *$p<0.05$.

**Supplementary Figure 2**

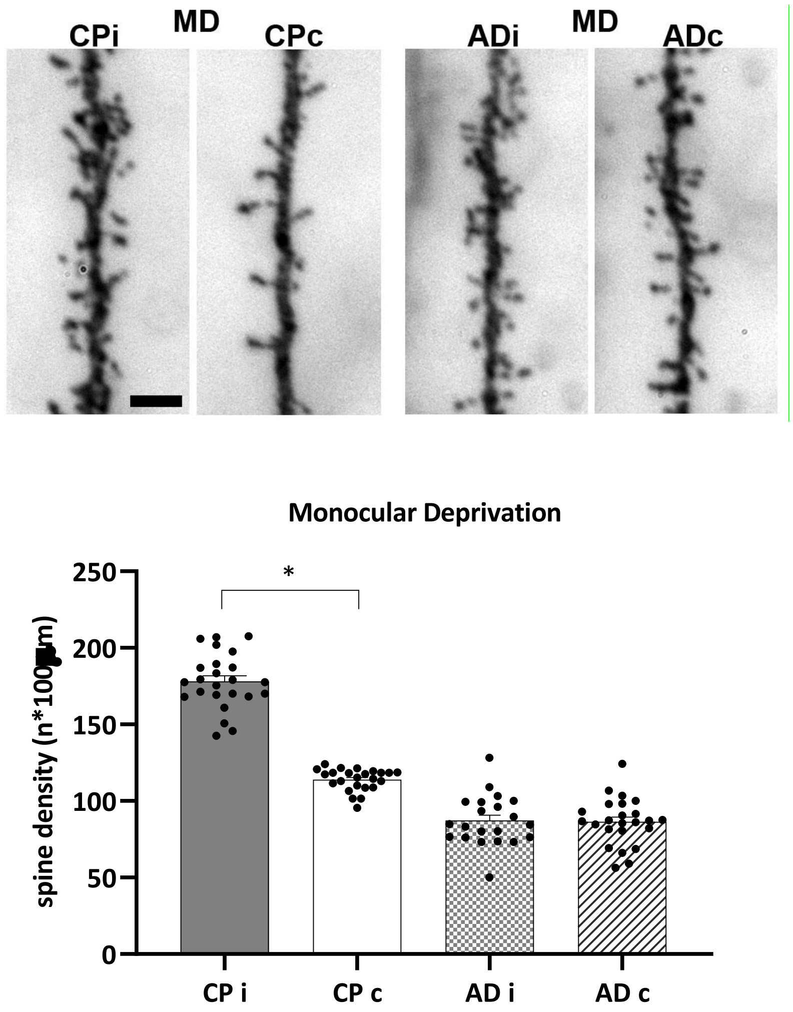


**Supplementary Figure 2:** Monocular deprivation reduces spine density in the contralateral visual cortex in critical period mice, but not in adult ones. Representative images of apical dendrites from the ispilateral and contralateral visual cortex of critical period (CPi and CPc) and adult (ADi and ADc) deprived (MD, right eye) mice. Bar: 5 µm. The scatter plot represents the spine density (n protrusions*100 µm) in the ispilateral (CPi) and contralateral (CPc) visual cortex of n=6 deprived CP mice, and in the ispilateral (ADi) and contralateral (ADc) visual cortex of n=6 deprived adult mice. Dots represent spine density values of individual dendrites; mean is marked as a bar±SEM. Mean spine density: CPi: 178.0±3.7 n=24 dendrites vs. CPc: 113.9±1.5 n=28 dendrites, n=6 mice, Student´s t-test, *p<0.05; ADi: 87.2±3.6, n=21 dendrites vs. ADc: 86.4±3.1 n=24 dendrites, n=6 mice, Student´s t-test, p=0.88.

**Supplementary Figure 3**

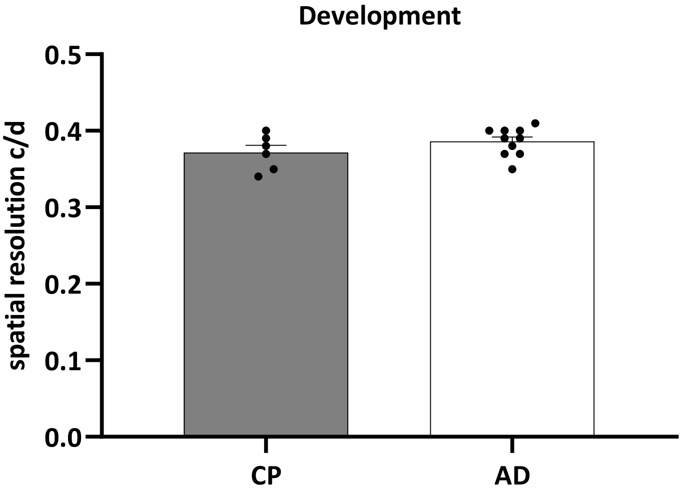


**Supplementary Figure 3:** Visual acuity is similar between critical period and adult mice. The scatter plot represents the spatial resolution (c/d) of n=6 critical period (CP) mice and n=10 adult (AD) mice. Visual acuity of the right and left eye were pooled. Dots represent VA values of individual mice; mean is marked as a bar±SEM. Mean VA: CP: 0.37±0.01 vs. AD: 0.38±0.01, Student´s t-test, p=0.85.

### Supplementary Figure 4

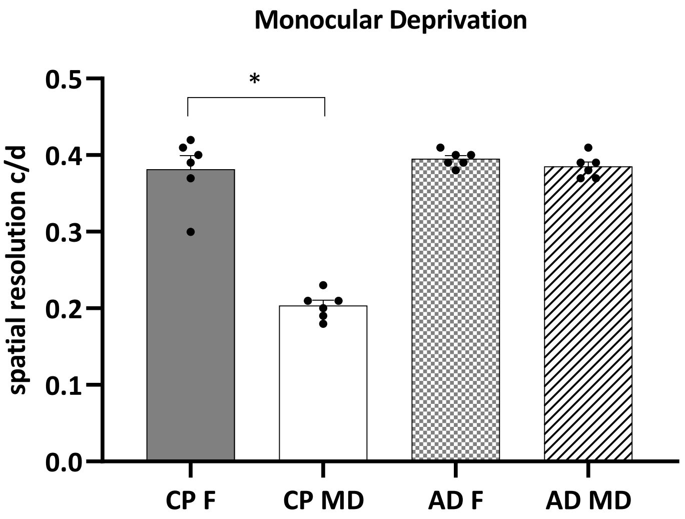


**Supplementary Figure 4:** Monocular deprivation reduces visual acuity through the deprived eye in critical period mice, but not in adult ones. The scatter plot represents the spatial resolution (c/d) through the fellow eye (CP F) and the deprived eye (CP MD) of n=6 critical period (CP) mice, and through the fellow eye (AD F) and the deprived eye (AD MD) of n=6 adult (AD) mice. Dots represent VA values of individual mice; mean is marked as a bar±SEM. Mean VA: CP F: 0.38±0.02 vs. CP F: 0.20±0.02, n=6, Student´s t-test, *p<0.05; AD F: 0.39±0.01 vs. AD MD: 0.38±0.01, n=6, Student´s t-test, p=0.81.
